# A quantitative fluorescence-based approach to study mitochondrial protein import

**DOI:** 10.1101/2022.10.10.511594

**Authors:** Naintara Jain, Ridhima Gomkale, Olaf Bernhard, Peter Rehling, Luis Daniel Cruz-Zaragoza

## Abstract

Mitochondria play central roles in cellular energy production and metabolism. Most of the proteins that are required to carry out these functions are synthesized in the cytosol and imported into mitochondria. A growing number of metabolic disorders arising from mitochondrial dysfunction can be traced to errors in mitochondrial protein import. The mechanisms underlying the import of precursor proteins are commonly studied by using radioactively-labeled precursor proteins, which are imported into purified mitochondria. Here, we establish a fluorescence-based import assay to analyze protein import into mitochondria. We show that fluorescently-labeled precursors enable import analysis with similar sensitivity to those using radioactive precursors, yet they provide the advantage of quantifying import with picomole resolution. We adapted the import assay to a 96-well plate format allowing for fast analysis in a screening-compatible format. Moreover, we show that fluorescently labeled precursors can be used to monitor the assembly of the F1F0 ATP-synthase in purified mitochondria. Thus, we provide a sensitive fluorescence-based import assay that enables quantitative and fast-import analysis.

## Introduction

Mitochondria play central roles in cellular metabolism and signaling processes (Nunnari & Suomalainen, 2012). While mitochondria possess a small genome, most mitochondrial proteins are nuclear encoded and imported after their synthesis in the cytosol (Pfanner *et al*, 2019; Richter-Dennerlein *et al*, 2015; Neupert & Herrmann, 2007; Araiso *et al*, 2022). The mitochondrial proteome comprises of more than 1,000 different proteins in the yeast *Saccharomyces cerevisiae* (Di Bartolomeo *et al*, 2020; Wiedemann & Pfanner, 2017; Morgenstern *et al*, 2017; Sickmann *et al*, 2003). The translocation of the precursor proteins across the outer and inner mitochondrial membranes requires multi-subunit protein translocation machineries in the membranes. The TOM (translocase of the outer membrane) complex facilitates translocation through the outer membrane while TIM (translocase of the inner membrane) complexes mediate translocation of precursors across the inner membrane (Lill & Neupert, 1996; Berthold *et al*, 1995; Wiedemann & Pfanner, 2017; Araiso *et al*, 2022). The majority of precursor proteins are directed across both membranes by N-terminal presequences, consisting of amphipathic alpha helices which are recognized by receptors in the TOM and TIM23 complex (Chacinska *et al*,2009; Geissler *et al*, 2002; Schulz *et al*, 2011; Yamano *et al*, 2007; Yamamoto *et al*,2009; Brix *et al*, 1997; Roise *et al*, 1986; Vögtle *et al*, 2009; Neupert & Herrmann, 2007; Araiso *et al*, 2022). The import across the TIM23 complex into the matrix requires membrane potential across the inner membrane (Δψ) and the activity of the presequence translocase - associated motor (PAM) complex. Upon import into the matrix, the presequence is cleaved by the mitochondrial processing peptidase (MPP) (Schulz *et al*, 2015; Wiedemann & Pfanner, 2017; Mossmann *et al*,2012).

Import into mitochondria is commonly studied by utilizing an *in vitro* import assay in which [^35^S]-labeled precursor proteins are imported post-translationally into isolated mitochondria (Harmey *et al*, 1977; Maccecchini *et al*, 1979). For this, radiolabeled precursors are synthesized in reticulocyte lysates and incubated with purified mitochondria. Dissipation of the membrane potential by inhibitors of the OXPHOS system and uncouplers is used to block import. A protease treatment of the reaction following import removes non-imported proteins from the system. Samples are analyzed by SDS- or BN-PAGE, and proteins are visualized by autoradiography. This *in vitro* import assay has been instrumental in dissecting the mechanisms of protein translocation across the mitochondrial membrane as it provides high sensitivity and kinetic resolution. However, absolute quantitative information on the imported amounts of precursors is difficult to obtain in this setup and the use of isotopes requires special safety precautions that are not readily available to all researchers. Moreover, the radioactive approach is difficult to combine with high-throughput screening approaches.

Here we report on a fluorescence-based method to monitor *in vitro* mitochondrial protein import using precursor-fluorophore fusion protein as a substrate. The non-radioactive method is sensitive, fast, and it allows to work with chemical quantities of import competent protein. The fluorescent approach provides the advantage of a fully quantitative output with picomolar resolution and the potential to perform import in a plate-format for rapid results. We show that, in addition to monitoring protein import, fluorescently-labeled proteins can also be utilized to analyze assembly of protein complexes in purified mitochondria.

## Results and discussion

### Jacl_488_ fluorescent precursor enables quantitative import analysis

Based on a previous observation, that a fluorescently labeled precursor protein retains the ability to be imported into mitochondria (Cruz-Zaragoza *et al*, 2021), we set out to establish a non-radioactive standard import assay. For the initial set of experiments, constructs consisting of the *S. cerevisiae* Jac1 protein with its authentic N-terminal presequence fused to a C-terminal FLAG tag was used (Fig 1A). The Jac1 used here carried a C145A exchange and an additional cysteine residue at the C-terminal allowing for addition of a fluorophore. For purification of the construct from *E. coli*, the protein carried a His-tag, which was cleaved off post-purification through a flanking SUMO protease site, preserving the N-terminus of the protein. After purification, the Jac1 fusion-protein was modified by maleimide-mediated addition of a DyLight fluorophore to the terminal cysteine residue (Jac1_488_). Next, we imported the precursor into isolated mitochondria. Samples were split after the import reaction and treated with Proteinase K (PK) to remove non-imported precursor. As a negative control, the membrane potential was dissipated prior to import. Samples were subjected to SDS-PAGE and gels scanned at the DyLight fluorophore emission range using a fluorescence scanner (Fig 1B). The Jac1_488_ precursor was imported into mitochondria in a time and membrane potential-dependent manner as apparent in the protease treated samples (Fig 1B). However, Jac1_488_ did not display efficient processing upon import. Based on results from import assays of various other protein constructs tested during this study, this is not a general phenomenon observed for fluorescent precursors, but more specific to certain proteins and presequences. After confirming that the substrate imported efficiently into mitochondria, we aimed to obtain quantitative data on the imported protein amounts. To this end, dilutions of the purified precursor protein were used as a standard and loaded together with the import samples on the gel (Fig 1C). A titration curve of the precursor standard was plotted to determine the absolute amount of imported protein per μg of mitochondria. We calculated that about 0.84 pmol protein was imported per μg mitochondria after 10 min and about 1.2 pmol after 15 min (Fig 1E). In this timeframe the import reaction was still in the linear range (Fig 1D). We concluded that fluorescently labeled precursors can be used for *in vitro* import and that the assay allowed us to obtain quantitative data on mitochondrial import. Accordingly, an absolute comparison between different substrates and import conditions can be obtained.

**Figure 1.**
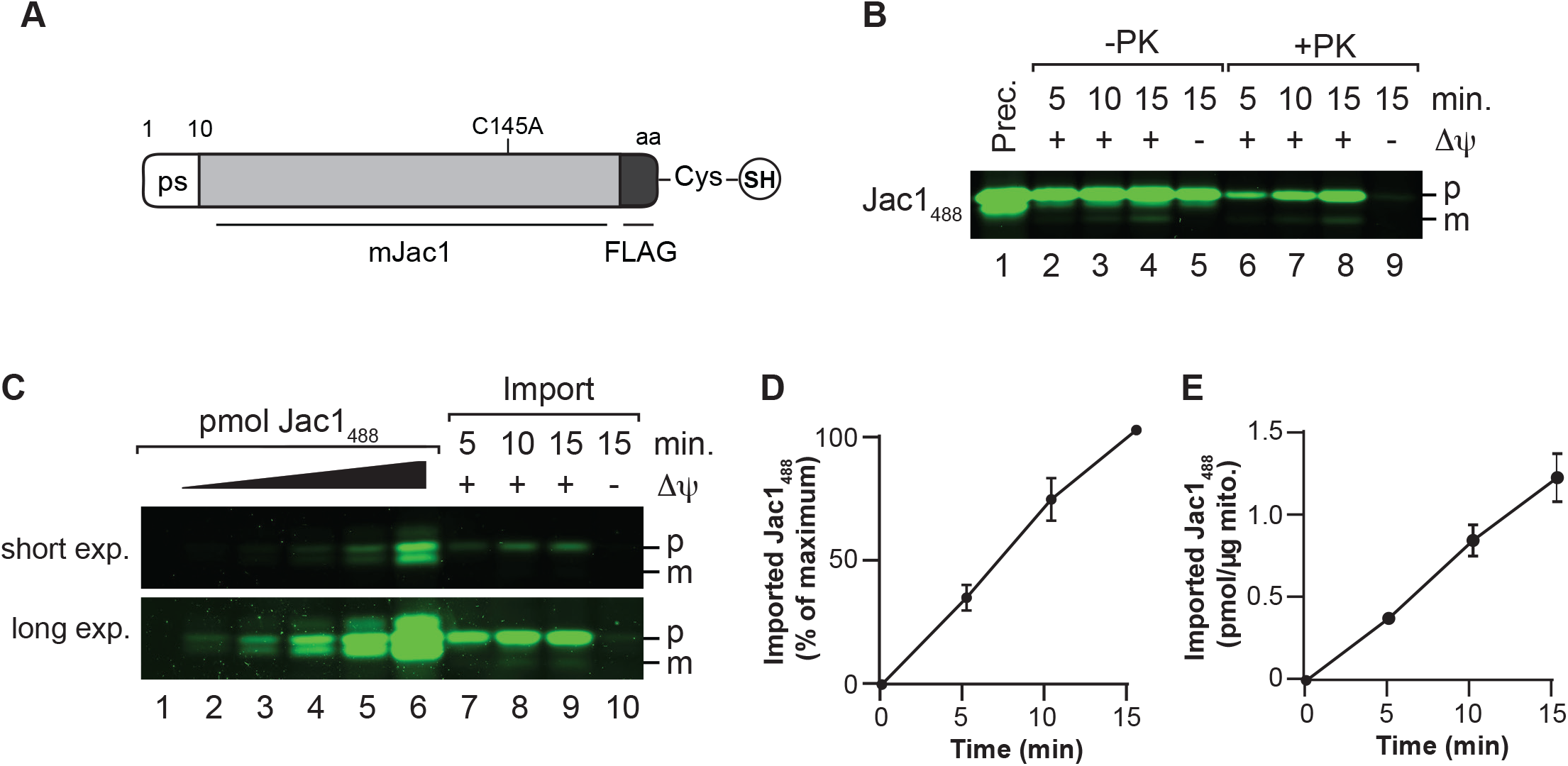
Import of fluorescent precursor into mitochondria. A. Schematic presentation of the modified Jac1 protein with C-terminal cysteine for fluorophore addition. B. Jac1_488_ was imported into purified mitochondria for indicated times and treated with Proteinase K (PK) or left untreated as indicated, Prec., purified precursor protein; p, precursor; m, mature protein; ΔΨ, membrane potential. C. Jac1_488_ import into mitochondria as in B, right side; Jac1_488_ protein dilution series, left side. D. Quantification of import of Jac1_488_ (maximal signal at 15 min import time, 100%); error bars indicate standard error of mean (SEM) (n=3). E. Quantification of absolute imported amounts in picomoles Jac1_488_ per μg of mitochondria; error bars indicate SEM (n=3).

### Jac1_488_ enables functional analysis of the import machinery

*In vitro* import into mitochondria is the key technology to dissect the mechanisms and components of protein translocation. To assess if the fluorescently-labeled precursor could be used to analyze defects in protein transport, we imported Jac1_488_ into purified yeast mitochondria with defects in the import machineries. Therefore, a temperature sensitive mutant of *Tim44 (tim44-804*) and a yeast strain in which the *TIM50* gene was under control of a GAL-promotor, allowing to decrease steady state levels of Tim50, were selected (Geissler *et al*, 2002; Hutu *et al*, 2008). Tim50 is the essential, central presequence receptor of the TIM23 complex and required for precursor import (Geissler *et al*, 2002; Yamamoto *et al*, 2002; Schulz *et al*, 2011; Qian *et al*, 2011; Mokranjac *et al*, 2003). Mitochondria depleted for Tim50 were isolated from *S. cerevisiae* (Schulz *et al*, 2011) and steady state protein levels were analyzed to confirm efficient knockdown. While the Tim50 levels were reduced in the mutant strain, Tom70, Tim23, and Hsp70 levels remained similar to the wild-type (WT), indicating that the remaining import machinery constituents were not affected by the knockdown (Fig 2A). Jac1_488_ was imported into WT and Tim50-depleted mitochondria (Fig 2B). As expected, Jac1_488_ import was severely affected in the mutant mitochondria, which amounted to about 40% of the WT import after 15 min (Fig 2C). The observed import defect matched the decrease in import observed with radioactively labeled Jac1 (Fig 2D & E).

**Figure 2.**
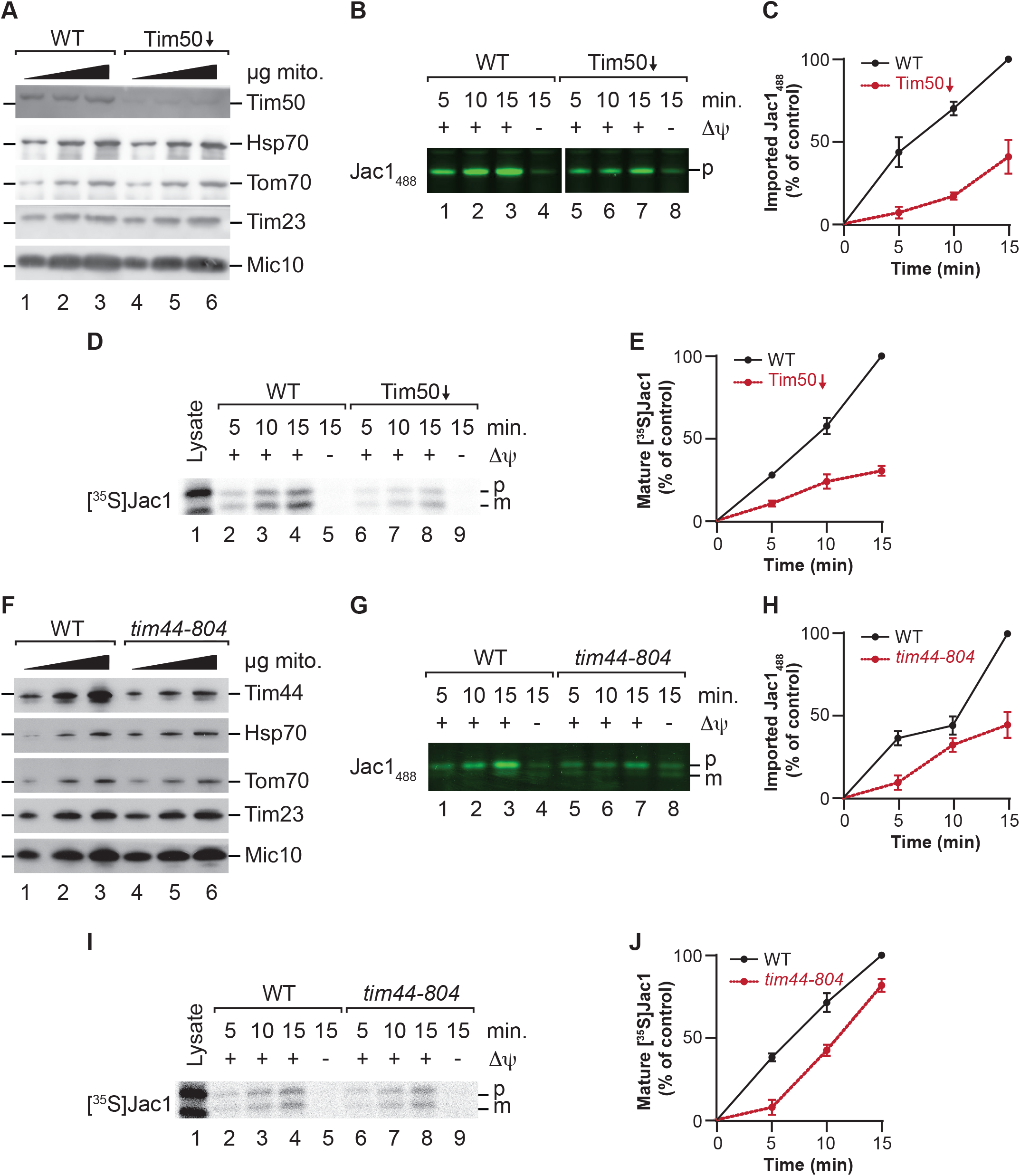
Import into mutant mitochondria. A. Steady state protein levels in wild-type (WT) and Tim50-depleted mitochondria visualized by Western blotting and immunodetection with indicated antisera. B. Jac1_488_ was imported into wild-type (WT) and Tim50-depleted mitochondria for indicated times and samples were treated with Proteinase K. p, precursor C. Quantification of Jac1_488_ import into wild-type (WT) and Tim50-depleted mitochondria. The amount of imported protease-protected protein in WT mitochondria at 15 min was set to 100%; error bars indicate SEM (n=3). D. [^35^S]Jac1 was imported into wild-type (WT) and Tim50-depleted mitochondria for indicated times and samples were treated with Proteinase K. p, precursor; m, mature. E. Quantification of [^35^S]Jac1 import into wild-type (WT) and Tim50-depleted mitochondria. The amount of imported protease-protected protein in WT mitochondria at 15 min was set to 100%; error bars indicate SEM (n=3). F. Steady state protein levels in wild-type (WT) and *tim44-804* mitochondria visualized by Western blotting and immunodetection with indicated antisera. G. Jac1_488_ was imported into wild-type (WT) and *tim44-804* mitochondria for indicated times and samples were treated with Proteinase K. p, precursor; m, mature. H. Quantification of Jac1_488_ import into wild-type (WT) and *tim44-804* mitochondria. The amount of imported protease-protected protein in WT mitochondria at 15 min was set to 100%; error bars indicate SEM (n=3). I. [^35^S]Jac1 was imported into wild-type (WT) and *tim44-804* mitochondria for indicated times and samples were treated with Proteinase K. p, precursor; m, mature. J. Quantification of [^35^S]Jac1 import into wild-type (WT) and *tim44-804* mitochondria. The amount of imported protease-protected protein in WT mitochondria at 15 min was set to 100%; error bars indicate SEM (n=3).

Tim44 is a constituent of the mitochondrial import motor and required for matrix protein transport (Schneider *et al*, 1994; Blom *et al*, 1993). We used a Tim44 temperature-conditional yeast mutant strain (*tim44-804*), which displays an import defect upon shift of mitochondria to 37°C (Hutu *et al*, 2008). Western blot analysis of the protein steady state levels in *tim44-804* displayed slightly reduced amounts of Tim44 in mitochondria while other analyzed translocase constituents were not decreased in the mutant (Fig 2F). Import of Jac1_488_ into WT and *tim44-804* mutant mitochondria showed a mutant-specific decrease in import (Fig 2G1 & H). It is interesting to note that the decrease in import was less pronounced when assayed using the radiolabeled Jac1 (Fig 2I & J). The increased import defect observed for the fluorescently-labeled precursor is possibly due to the larger quantities of precursor applied to mitochondria compared to the radiolabeled counterpart, which challenges the import machinery for translocation. It is also conceivable that chaperones that are associated to the radiolabeled precursor after synthesis in the reticulocyte lysate may stimulate the import by unfolding the precursor for import. In summary, these results confirm that import defects can be efficiently assayed using the fluorescently-labeled precursor.

### Effect of presequences swapping on import efficiency

As a way to utilize this method to study different characteristics of a precursor, we synthesized Jac1 constructs with different targeting signals. For this, the authentic presequence of Jac1 was replaced with presequences of Idh1 (Isocitrate dehydrogenase 1) or Aco1 (Aconitase 1). These presequences were selected due to their similarity to the Jac1 presequence regarding length and charge (^1-10^Jac1^ps^, +2.26 net charge; ^1-12^Idh1^ps^, +2,27 net charge; ^1-16^Aco1^ps^, +3,27 net charge) (Fig 3A). The same procedure was followed for purification and modification of these constructs as described above for the authentic Jac1. All constructs were modified with three different DyLight fluorophores (with excitation wavelengths 488, 680, and 800 nm), with the assumption that the modification should not affect import but allow to multiplex import using precursors with different fluorophores in the same sample (Fig 3B). Subsequently, all precursor constructs were imported into purified mitochondria using the Jac1_488_ precursor as a control. After import, samples were analyzed by SDS-PAGE and the fluorescence signal of the imported proteins was quantified. All fusion proteins displayed membrane potential-dependent import that increased with time (Fig 3C & 3E). Despite a slightly more positive charged and longer presequence in case of the Aco1, the import of the pAmJac1 (mature Jac1 protein with Aco1 presequence) showed no significant difference in import compared to the Jac1 control (Fig 3D). Similar to Jac1, the pAmJac1 variants did not display efficient processing after import (Fig 3C). In the import experiments, pImJac1 (mature Jac1 protein with Idh1 presequence) precursor variants differed slightly compared to the control, despite the almost identical size and charge of the presequence (Fig 3F). However, upon import of pImJac1, we observed efficient processing of the Idh1 presequence (Fig 3E). These two examples showed that indeed the import of fluorescently labeled precursors represents a means to analyze precursor properties for mitochondrial protein import. In addition, in both cases, the import of the constructs was not affected by the choice of fluorophore (Fig 3D & 3F). Accordingly, different combinations of fluorophores can be used to monitor import. Since there was no bleed-through of the fluorescence between different scanning channels, the use of differently labeled precursors will enable multiplexing of import assays using different precursors, tagged by different fluorophores.

**Figure 3.**
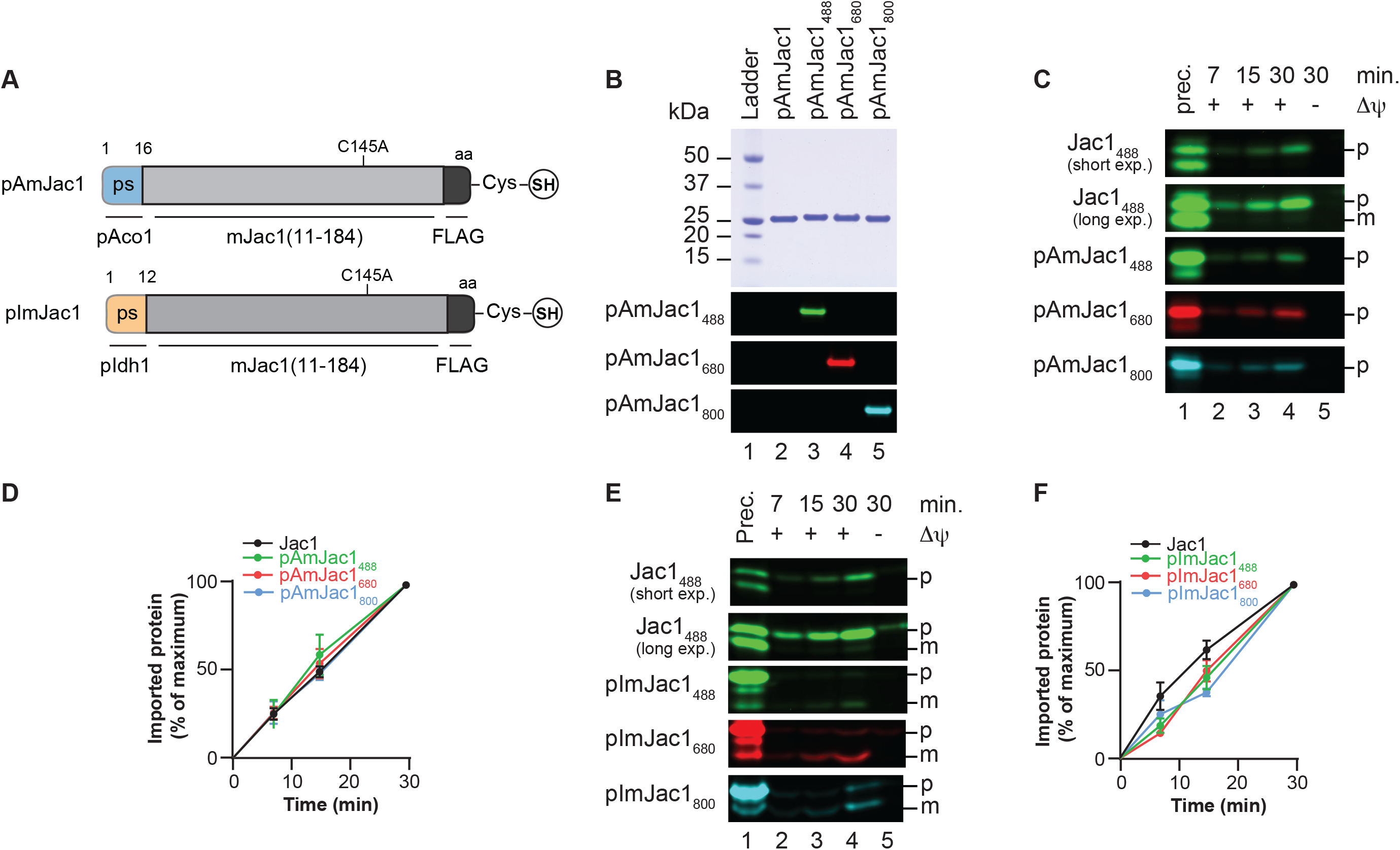
Import of precursor variants into mitochondria. A. Schematic presentation of pAmJac1 and pImJac1, with presequences derived from Aco1 and Idh1, respectively, fused to the N-terminus of the mature Jac1 portion. B. Purified pAmJac1 was conjugated with different fluorophores as indicated. Detected without bleed through under fluorescence imaging conditions. C. Jac1_488_ and pAmJac1 conjugated to DyLight 488, 680, and 800 were imported into wild-type mitochondria for indicated times and samples were treated with Proteinase K. Prec., purified precursor protein; p, precursor; m, mature. D. Quantification Jac1_488_ and pAmJac1 conjugated to DyLight 488, 680, and 800 import into wild-type (WT). The amount of imported protease-protected samples at 30 min was set to 100%; error bars indicate SEM (n=3). E. Jac1_488_ and pImJac1 conjugated to DyLight 488, 680 and 800 were imported into wild-type mitochondria for indicated times and samples treated with Proteinase K. Prec., purified precursor protein; p, precursor; m, mature F. Quantification Jac1_488_ and pImJac1 conjugated to DyLight 488, 680 and 800 import into wild-type (WT). The amount of imported protease-protected samples at 30 min was set to 100%; error bars indicate SEM (n=3).

### Assessing import in 96-well plate format

While the current standard import assay requires separation of proteins by PAGE analysis to observe the labeled protein, we were curious if we could adapt the process to a plate format in which samples could be analyzed rapidly with a fluorescence plate reader. Therefore, we performed the import assay using the pAmJac1_680_ precursor as described above and subsequently treated the mitochondria with proteinase K. Following import, mitochondria were re-isolated, resuspended, and transferred to 96-well plates (Fig 4A). For comparison, the import reactions were normally split and analyzed in a plate assay and by SDS-PAGE. To enable quantification, a dilution series of the precursor was measured as a standard (Fig 4B). Fluorescence measurements of pAmJac1_680_ import reactions revealed a time-dependent localization of the construct to mitochondria (Fig 4C). The fluorescence measured in the membrane potential depleted sample was subtracted from the individual measurements to correct for background binding. A quantification of the plate assay showed that 0.06, 0.12, and 0.2 pmol of pAmJac1_680_, protein were imported per μg mitochondria at 5, 10, and 15 minute time points respectively (Fig 4D). Accordingly, a plate-format is suitable to assess import of labeled precursors, which can be quantified much faster compared to a gel-based system. A comparison between the results of the plate format and the standard PAGE analysis showed that both types of analysis provided similar data on the kinetics of the import reaction (Fig 4E). A quantitative comparison showed only a small divergence between the two approaches that can possibly be attributed to precursor fragments that are detected by total fluorescence but not resolved on the gel.

**Figure 4.**
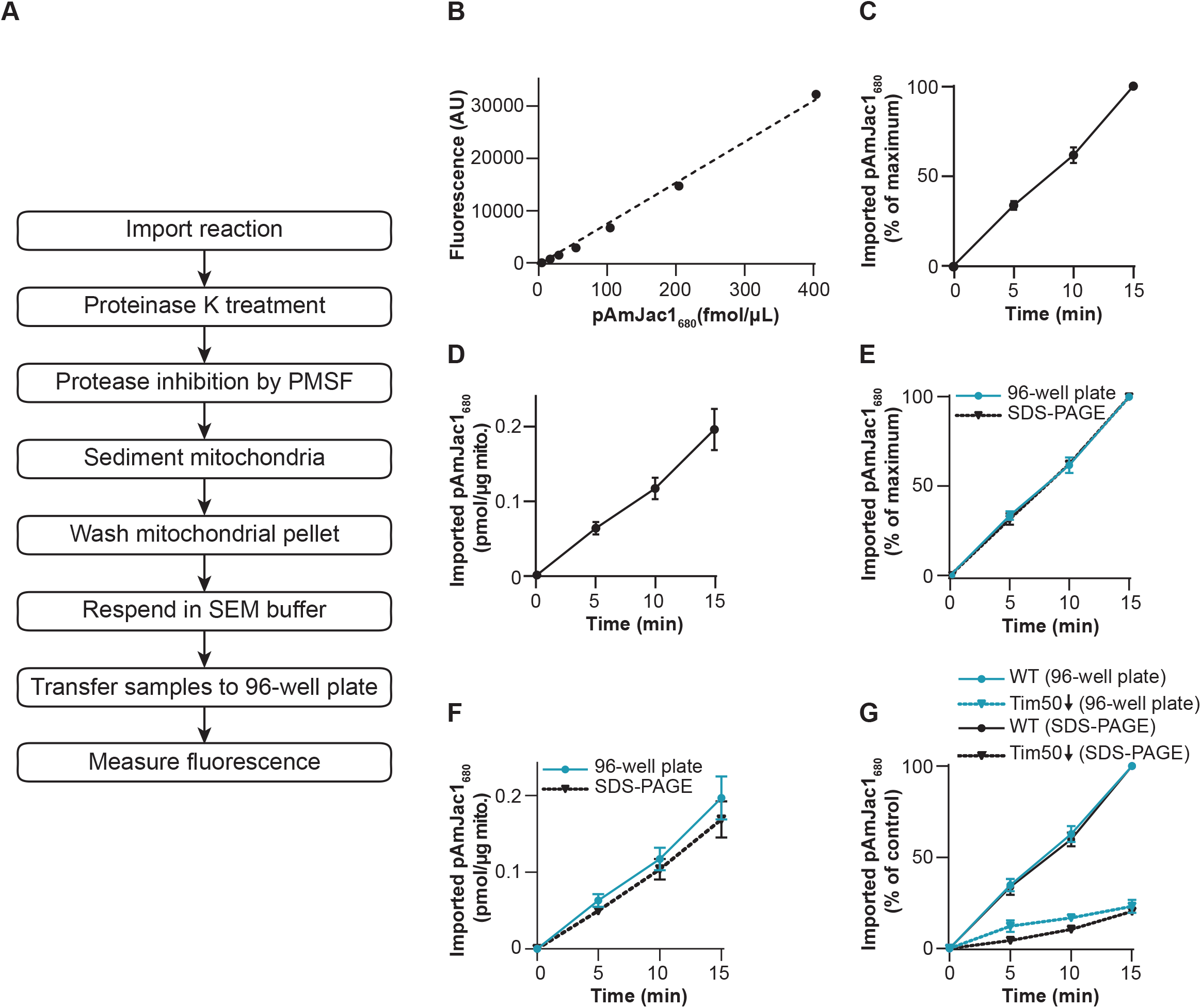
Analyzing import in a multi well plate format. A. Workflow for transfer of imported samples into 96-well plate for readout. B. Standard curve depicting fluorescence signal plotted against concentration of pAmJac1_680_ dilutions. C. Import of pAmJac1_680_ into wild-type mitochondria. After import and proteinase K treatment fluorescence was measured in 96-well format. The amount of imported protease-protected samples at 15 min was set to 100%; error bars indicate SEM (n=8). D. Quantification of picomoles of pAmJac1_680_ imported per μg mitochondria as assessed in 96-well format; error bars indicate SEM (n=8). E. pAmJac1_680_ was imported into purified mitochondria and samples analyzed by SDS-PAGE or in 96-well format. The amount of imported, protease-protected samples at 15 min was set to 100% in each case; error bars indicate SEM (n=8 for 96-well plate analysis, n=3 for SDS-PAGE). F. Comparison of absolute import of pAmJac1_680_ into wild-type mitochondria quantified from 96-well format and SDS-PAGE. Error bars indicate SEM (n=8 for 96-well plate analysis, n=3 for SDS-PAGE). G. pAmJac1_680_ was imported into wild-type (WT) and Tim50-depleted mitochondria for indicated times and samples treated with proteinase K. Samples were analyzed in 96-well format and by SDS-PAGE. The amount of imported protease-protected protein in WT mitochondria at 15 min was set to 100%; error bars indicate SEM (n=5 for 96-well plate analysis, n=4 for SDS-PAGE).

Next, we addressed if the plate format could be applied to analyze defects in mitochondrial import. For this we performed import into wild-type and Tim50-depleted mitochondria (Fig 4G). Similar import kinetics were apparent with both approaches and the magnitude of the mutant import defect was comparable. Accordingly, the analysis of import reactions by plate assay provided an efficient means for import studies. Yet, while the plate-format saves time and provides accurate readouts on import kinetics and efficiency, it cannot provide information on protein processing after import and can only give a value for the total imported, protease protected protein amounts.

### Fluorescent Atp5 enables analysis of complex assembly

During the course of the study, we tested various protein-fluorophore fusion constructs. Among these was Atp5_488_, a subunit of the F_1_F_o_ ATP-synthase, which displayed efficient processing comparable to that seen in the autoradiograph of import reactions using a radioactive Atp5 version (Fig 5A). The low background and strong signal suggested that it could be a good candidate for the 96-well plate import readout. Hence, we used this precursor to perform further import experiments and to corroborate the results with import of radioactive Atp5. We tested the dependence of Atp5 on the mitochondrial membrane potential for import using the uncoupler CCCP (carbonyl cyanide-m-chlorophenylhydrazone) that allows the dissipation of the ΔΨ in a concentration dependent manner (Martin *et al*, 1991). To this end, we titrated increasing amounts of CCCP and analyzed protein import into mitochondria. We observed that the amount of mature Atp5_488_ decreased with increasing CCCP concentration (Fig 5B). We compared the decrease of import with increasing CCCP concentrations between the fluorescent Atp5_488_ protein quantified using a gel system and a 96-well plate format (Fig 5B & C) and [^35^S]Atp5 analyzed by autoradiography (Fig 5D & E). In all three cases a similar ΔΨ-dependence of the import was observed supporting the usefulness of the fluorescence approach for analyzing the bioenergetics of protein import.

**Figure 5.**
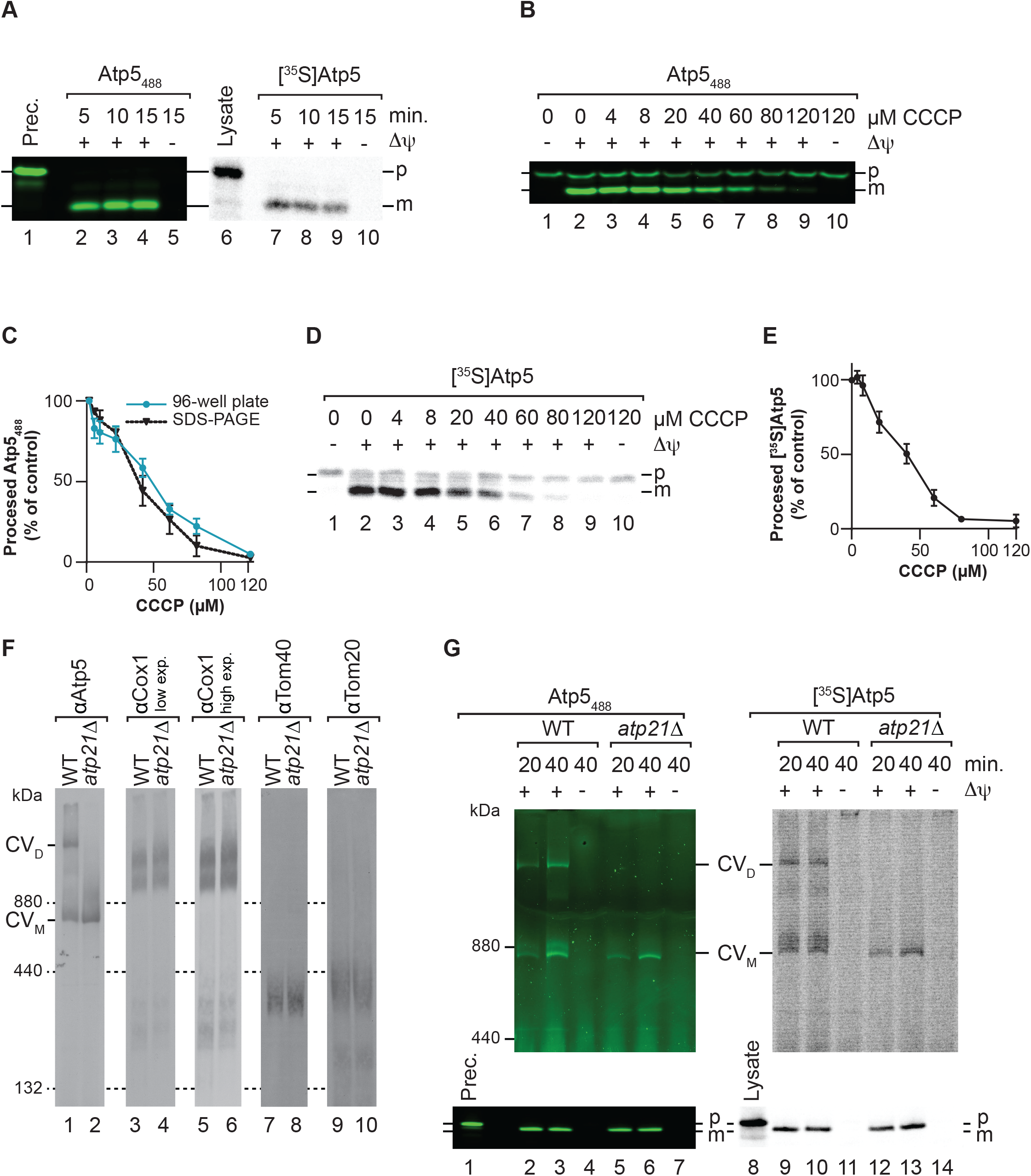
Fluorescence based protein assembly. A. Atp5_488_ (left panel) or [^35^S]Atp5 (right panel) were imported into mitochondria for indicated times and in the presence or absence of a membrane potential (ΔΨ). After proteinase K treatment, samples were separated by SDS-Page and analyzed by fluorescence scanning or digital autoradiography. Prec., purified precursor protein; p, precursor; m, mature protein. B. Import of Atp5488 with increasing concentrations of CCCP to study dependency of import on membrane potential (ΔΨ). P, precursor; m, mature protein. C. Comparison of import of Atp5_488_ with increasing CCCP concentrations quantified from 96-well format and SDS-PAGE. The amount of imported protease-protected samples in the absence of CCCP was set to 100%; error bars indicate SEM (n=4 for 96-well plate analysis, n=2 for SDS-PAGE). D. [^35^S]Atp5 was imported with increasing CCCP concentrations as described in B. Samples were separated by SDS-PAGE and analyzed by digital autoradiography. E. Quantification of import of [^35^S]Atp5 with increasing CCCP concentrations. The amount of imported protease-protected protein in the absence of CCCP was set to 100%; error bars indicate SEM (n=3). F. BN-PAGE steady state analysis of complex V, complex IV, and TOM in mitochondria isolated from WT and *atp21*Δ yeast strains. G. Atp5_488_ (left panel) or [^35^S]Atp5 (right panel) were imported into wild-type and *atp21*Δ mitochondria for indicated times and in the presence or absence of a membrane potential (ΔΨ). After proteinase K treatment, samples were solubilized in digitonin buffer and separated by BN-Page. Aliquots of the samples was analyzed by SDS-PAGE (lower panels). Subsequently proteins were visualized by fluorescence scanning or digital autoradiography. Complex V dimer, (CV_D_); monomeric (CV_M_).Prec., purified precursor; p, precursor; m, mature protein.

The availability of a subunit of an OXPHOS complex suitable for protein import led us to ask if a fluorescence import approach was appropriate for the analysis of OXPHOS complex assembly. To address this, we purified mitochondria from an *atp21Δ* strain which lacks the ability to form complex V dimers, as can be analyzed by Blue Native PAGE (Fig 5F, lanes 1-2). Radiolabeled Atp5 and Atp5_488_ efficiently assembled into monomeric and dimeric ATP synthase in wild-type mitochondria in a membrane potential-dependent manner. In *atp21Δ* mutant mitochondria both proteins only assembled into the monomeric ATP synthase as these mitochondria lack the dimeric form (Fig 5G). Accordingly, OXPHOS protein complex assembly can be studies with a fluorescent fusion protein. The non-radioactive approach provides similar information and sensitivity as the radioactive assay.

## Conclusion

Here, we present a new approach to study protein import into mitochondria. The commonly used method in the field is the synthesis of precursor proteins by *in vitro* translation using radioactive labeling for detection after protein or protein complex separation by PAGE analysis. Yet, the use of radioactive substances is problematic for many labs due to safety considerations related to the technique. Different approaches have been previously attempted to address protein import with non-radioactive strategies. In principle the use of recombinant proteins is established and has the advantage of providing import substrates in chemical and translocase saturating quantities with western blot-based detection (Voos et al., 1993; Mokranjac et al., 2005; Schulz & Rehling, 2014). In a recent study, Pereira et al. (2019) established an approach utilizing a bipartite luminescence system that employs NanoBiT - a split luciferase - to monitor protein import. Yet, this approach requires the introduction of one of the components in advance of the import assay into mitochondria making it less flexible to use in a mutant context. Here, we provide a non-radioactive alternative that is based on the use of fluorescently-labeled proteins and that is complementary to the existing strategies, filling the analytic gaps of the existing techniques. Chemical modification with fluorescent dyes allows the use of gel based and multi-well plate-based approaches. Moreover, as the absolute amounts of precursors can be determined, it is possible to easily assess the protein import in quantitative terms. Finally, the in-gel and in 96-well format offer a faster detection mode than radioactive assay. Beyond these applications, we show that after the fluorescently-labeled protein import, it can assemble into its target complex. Hence, the *in vitro* import system allows analysis of assembly processes in similar manner as with radiolabeled proteins and broadens the scope of non-radioactive approaches.

## Material and methods

### Plasmid generation for protein expression in bacteria

For the production of recombinant Jac1-FLAG-Cys, a plasmid reported previously was used (Cruz-Zaragoza *et al* 2021) where Jac1 is tagged at the N-terminus with a His14-SUMO-tag. To swap the Jac1 presequence, the sequence encoding presequences of *Saccharomyces cerevisiae* Aco1 and Idh1 were added by PCR at the 5’ of Jac1 ORF, removing the Jac1 presequence, followed by Gibson assembly. *ATP5* gene was amplified from *S.cerevisiae* genomic DNA and ligated into pJET vector using CloneJET PCR Cloning kit (Thermo Scientific). Site-directed mutagenesis was used to recode the cysteine at position 117 to alanine. The expression vector was amplified by PCR while removing Jac1 ORF but preserving FLAG-tag and C-terminal cysteine coding sequence. Finally, the mutant ORF was amplified by PCR and inserted in frame after the His14-SUMO-tag sequence from the amplified vector to generate a plasmid for the expression of His14-SUMO-Atp5^C117A^-FLAG-Cys.

### Recombinant protein expression and purification

Recombinant Jac1, its variants with different presequences (pAmJac1 and pImJac1) and Atp5^C117A^ were purified as follows. Plasmids were transformed into *Escherichia coli* BL21 Tuner (DE3) strain (Sigma-Aldrich) or Rossetta DE3 (Novagen) in case of Atp5^C117A^. One colony was inoculated into LB medium supplemented with 2% glucose and 50 μg/mL kanamycin. The preculture was incubated for eight hours at 30°C. The OD_600nm_ was determined and fresh medium was inoculated at initial OD_600nm_=0.1 and incubated overnight at 37°C. Next, the preculture was diluted in LB medium supplemented with 50 μg/mL kanamycin to OD_600nm_=0.05. The culture was incubated at 37°C until the OD_600nm_ reached 0.6-0.8. Protein expression was induced with 0.2 mM IPTG and incubated for five hours. Cells were harvested and kept at −80°C. Cells were resuspended in lysis buffer (40 mM Tris/HCl, 500 mM NaCl, 10 mM Imidazole, 1 mM PMSF, 0.2 mg/mL DNase1, 1x complete protease inhibitor cocktail (Roche), pH 7.4). Cell disruption was performed with an EmulsiFlex-C3 (AVESTIN). The lysate was cleared by centrifugation in an SS-34 rotor at 23,000 x g at 4°C for 60 minutes. Next, the supernatant was collected and injected in two HisTrap colums (Cytiva) in tandem pre-equilibrated in buffer A1 (40 mM Tris/HCl, 500 mM NaCl, 10 mM Imidazole, pH 7.4). After exhaustive washing with buffer A2 (40 mM Tris/HCl, 500 mM NaCl, 30 mM Imidazole, pH 7.4). Bound protein was eluted in a gradient 0-100% of buffer B (40 mM Tris/HCl, 500 mM NaCl, 500 mM Imidazole, pH 7.4). The fractions containing the protein of interest were pooled, and the buffer was exchanged with Desalting buffer (20 mM Tris/HCl, 150 mM NaCl, pH 7.4) in HiPrep^™^ 26/10 Desalting column (Cytiva). Protein concentration was determined and an adequate amount of His_6_-SUMO protease was added. The digestion was performed overnight at 4°C in the presence of 1 mM DTT and 5% of glycerol. Digestion efficiency was confirmed by SDS-PAGE. Next, imidazole 1 M pH 8.0 was added to the digestion mix to a final concentration of 20 mM. To deplete His_14_-SUMO-tag and the His6-tagged SUMO protease, an appropriate volume of Protino^®^ NiNTA-Agarose (MACHEREY-NAGEL) slurry was washed with Desalting buffer. Then, the digestion mix was added to the sedimented beads and incubated overnight at 4°C with end-to-end mixing. The unbound fraction was collected by gravity flow, and the bound protein was eluted with equivalent volume of buffer B. The quality of depletion was assessed by SDS-PAGE.

### Synthesis of protein-DyLight fluorescent adducts

Purified proteins were reduced with TCEP (Sigma-Aldrich). Excess TCEP was removed by buffer exchange to Maleimide buffer (100 mM potassium phosphate, 150 mM NaCl, 250 mM sucrose, 1mM EDTA, pH 6.6) in HiPrep^™^ 26/10 Desalting column (Cytiva). To fluorescently label the proteins, reduced protein was mixed with a 3:1 molar excess of DyLight_488_/DyLight_680_/DyLight_800_-maleimide (in dimethylformamide) and incubated overnight at 4°C. Unreacted maleimide groups were quenched with a 50-fold molar excess of cysteine. Finally, the buffer was exchanged to 20 mM HEPES, 150 mM KCl, 5% glycerol, pH 7.4 in HiPrep™ 26/10 Desalting column (Cytiva). The protein concentration was determined and set to 0.5-1 mg/mL. Aliquots of the labeled protein were kept at −80°C until used.

### Mitochondrial isolation

Yeast mitochondria were isolated using differential centrifugation (Meisinger et al., 2006). YP Media (1% yeast extract, 2% peptone) containing 2% glucose (YPD for primary cultures) or 3% glycerol (YPG for secondary cultures) was used as carbon source to grow wild-type YPH499 (MATa *ade2-101, his3*-Δ*200, leu2*-Δ*1, ura3-52, trp1*-Δ*63, lys2-801*), BY4741 wild-type (MATa *his3*Δ*1, leu2*Δ*0, met15*Δ*0, ura3*Δ*0*) and BY4741 *atp21D* yeast strains (Euroscarf) at 30°C with shaking till OD_600_ reached 1.5-2.5, after which they were harvested. The pellet was washed with water to remove remaining media and treated with DTT buffer (10 mM DTT, 100 mM Tris/HCl, pH 9.4) for 30 min at 30°C with shaking. Cells were washed with 1.2 M Sorbitol and treated with Zymolase buffer (20 mM KPO_4_, pH 7.4, 1.2 M sorbitol, and 0.57 mg/L zymolase) for 1 h at 30°C with shaking. Spheroplasts were harvested and resuspended in cold homogenization buffer (600 mM sorbitol, 10 mM Tris/HCl, pH 7.4, 1 g/L BSA, 1 mM PMSF, and 1 mM EDTA) before lysis using a homogenizer. Mitochondria were subsequently isolated using differential centrifugation and resuspended in SEM buffer. Protein concentration in the isolated mitochondria was determined using a Bradford assay and the final concentration adjusted to 10 mg/ml using SEM (250 mM sucrose, 20 mM MOPS/KOH pH 7.2, 1 mM EDTA) buffer before aliquoting and snap-freezing for storage at −80°C.

Mutant strain *tim44-804* (*MATa, ade2-101, his3*-Δ*200, leu2*-Δ*1, ura3-52, trp1*-Δ*63, lys2-801, tim44::ADE2*, [*pBG-TIM44-0804*]) and the corresponding wild-type cells were grown under permissive conditions, and mitochondria isolated as described previously (Hutu *et al* 2008).

Mitochondria depleted in Tim50, the AG55Gal strain (*MAT*a*, ade2-101, his3-Δ200, leu2-Δ1, ura3-52, trp1-Δ63, lys2-801, tim50::HIS3-PGAL1-TIM50*) and the corresponding wild-type cells were grown and processed as described previously (Schulz *et al*, 2011).

### Import into isolated mitochondria

For the synthesis of [^35^S] labeled Jac1 and Atp5 precursors, mRNA was generated using the mMessageMachine SP6 transcription kit (Invitrogen), following the manufacturer’s instructions. The mRNA obtained was used for *in vitro* translation in Flexi Rabbit Reticulocyte Lysate System (Promega) and the resulting lysate was directly used in the import reaction. The fluorescent precursors, as described above, were thawed just prior to import.

Import reactions were performed as described by (Ryan et al., 2001). Mitochondria were resuspended in import buffer (250 mM sucrose, 10 mM MOPS/KOH pH 7.2, 80 mM KCl, 2 mM KH_2_PO_4_, 5 mM MgCl_2_, 5 mM methionine, and 3% fatty acid-free BSA) and supplemented with 2 mM ATP and 2 mM NADH (as well as 5 mM Creatine phosphate and 1μg/μl Creatine kinase for experiments where the time for import exceeded 15 minutes). Import was performed by incubating samples at 25°C and stopped by the addition of 1% AVO (final concentration 1 μM valinomyin, 8 μM antimycin A and 20 μM oligomycin). The import samples were treated with 20 μg/ml proteinase K (PK) for 10 min on ice. Samples were treated with 2 mM PMSF, incubated on ice for 10 min, and centrifuged to sediment the mitochondria which were washed with SEM buffer and then analyzed by SDS-PAGE followed by western blotting and digital autoradiography (Amersham Typhoon, Cytiva) or fluorescent scanning (Starion FLA-9000, FujiFilm), based on the imported precursor. Signals were quantified using ImageQuant LT (GE Healthcare) with a rolling ball background quantification. Alternatively, for the 96-well format, the mitochondria were centrifuged after proteinase K and PMSF treatment, washed with SEM, resuspended in SEM buffer for transfer to a 96-well plate, and read on the Spark Multimode Microplate Reader (Tecan). Values of import were normalized by subtracting the signal from the AVO control sample and graphically represented using GraphPad Prism 8.

Mitochondria isolated from *tim44* temperature-conditional yeast mutant were incubated at 37°C for 15 minutes in import buffer prior to the addition of ATP, NADH and the precursor.

### Quantification of imported protein

Purified proteins were diluted in desalting buffer. A standard curve was plotted upon fluorescence measurement. This standard curve was used to calculate the corresponding total protein amount in the import samples.

### Membrane potential reduction – CCCP titration

Increasing amounts of the uncoupler CCCP were titrated into the import reaction to reduce the membrane potential (van der Laan et al., 2006). Import buffer used for the import reaction (as described above) was supplemented with 1% fatty acid-free BSA and 20 μM oligomycin. Mitochondria were kept at 25°C for 5 min prior to precursor addition.

## Acknowledgements

Funded by the Deutsche Forschungsgemeinschaft (DFG, German Research Foundation) under Germany’s Excellence Strategy - EXC 2067/1-390729940; SFB1190 (projects P13, PR); SFB860 (B01, PR); the Max Planck Society (PR), and the PhD program Molecular Biology - International Max Planck Research School and the Göttingen Graduate School for Neurosciences and Molecular Biosciences (GGNB; DFG grant GSC 226/1) (RG, NJ).

## Author Contributions

NJ, RG, PR and LDCZ developed the concept and design of the study. NJ, RG, OB and LDCZ performed the experiments and analyzed the data. NJ, PR, and LDCZ wrote the original draft. NJ, RG, PR, and LDCZ reviewed and edited the final draft of the manuscript. PR and LDCZ provided supervision.

## References

Araiso Y, Imai K & Endo T (2022) Role of the TOM Complex in Protein Import into Mitochondria: Structural Views. Annu Rev Biochem 91

Di Bartolomeo F, Malina C, Campbell K, Mormino M, Fuchs J, Vorontsov E, Gustafsson CM & Nielsen J (2020) Absolute yeast mitochondrial proteome quantification reveals trade-off between biosynthesis and energy generation during diauxic shift. Proc Natl Acad Sci U S A 117: 7524–7535

Berthold J, Bauer MF, Schneider HC, Klaus C, Dietmeier K, Neupert W & Brunner M (1995) The MIM complex mediates preprotein translocation across the mitochondrial inner membrane and couples it to the mt-Hsp70/ATP driving system. Cell 81: 1085–1093

Blom J, Kubrich M, Rassow J, Voos W, Dekker PJT, Maarse AC, Meijer M, Pfanner2 N, Maarse JC, Blom LA, et al (1993) The essential yeast protein MIM44 (encoded by MPI1) is involved in an early step of preprotein translocation across the mitochondrial inner membrane. Mol Cell Biol 13: 7364–7371

Brix J, Dietmeier K & Pfanner N (1997) Differential Recognition of Preproteins by the Purified Cytosolic Domains of the Mitochondrial Import Receptors Tom20, Tom22, and Tom70*.

Chacinska A, Koehler CM, Milenkovic D, Lithgow T & Pfanner N (2009) Importing Mitochondrial Proteins: Machineries and Mechanisms. Cell 138: 628

CHurt E, Muller U & Schatz Biocenter G (1985) The first twelve amino acids of a yeast mitochondrial outer membrane protein can direct a nuclear-encoded cytochrome oxidase subunit to the mitochondrial inner membrane. EMBO J 4: 3509–3518

Cruz-Zaragoza LD, Dennerlein S, Linden A, Yousefi R, Lavdovskaia E, Aich A, Falk RR, Gomkale R, Schöndorf T, Bohnsack MT, et al (2021) An in vitro system to silence mitochondrial gene expression. Cell 184: 5824–5837.e15

Geissler A, Chacinska A, Truscott KN, Wiedemann N, Brandner K, Sickmann A, Meyer HE, Meisinger C, Pfanner N & Rehling P (2002) The Mitochondrial Presequence Translocase: An Essential Role of Tim50 in Directing Preproteins to the Import Channel. Cell 111: 507–518

Harmey MA, Hallermayer G, Korb H & Neupert W (1977) Transport of cytoplasmically synthesized proteins into the mitochondria in a cell free system from Neurospora crassa. Eur J Biochem 81: 533–544

Hutu DP, Guiard B, Chacinska A, Becker D, Pfanner N, Rehling P & Van Der Laan M (2008) Mitochondrial protein import motor: Differential role of Tim44 in the recruitment of Pam17 and J-complex to the presequence translocase. Mol Biol Cell 19: 2642–2649

van der Laan M, Wiedemann N, Mick DU, Guiard B, Rehling P & Pfanner N (2006) A Role for Tim21 in Membrane-Potential-Dependent Preprotein Sorting in Mitochondria. Curr Biol 16: 2271–2276

Lill R & Neupert W (1996) Mechanisms of protein import across the mitochondrial outer membrane. Trends Cell Biol 6: 56–61

Maccecchini ML, Rudin Y, Blobel G & Schatz G (1979) Import of proteins into mitochondria: precursor forms of the extramitochondrially made F1-ATPase subunits in yeast. Proc Natl Acad Sci U S A 76: 343–347

Martin J, Mahlke K & Pfanners N (1991) Role of an energized inner membrane in mitochondrial protein import. Delta psi drives the movement of presequences. rNE J Biol Chem cc) 266: 18051–18057

Meisinger C, Pfanner N & Truscott KN (2006) Isolation of yeast mitochondria. Methods Mol Biol 313: 33–39

Mokranjac D, Paschen SA, Kozany C, Prokisch H, Hoppins SC, Nargang FE, Neupert W & Hell K (2003) Tim50, a novel component of the TIM23 preprotein translocase of mitochondria. EMBO J 22: 816–825

Mokranjac D, Sichting M, Popov-Čeleketić D, Berg A, Hell K & Neupert W (2005) The Import Motor of the Yeast Mitochondrial TIM23 Preprotein Translocase Contains Two Different J Proteins, Tim14 and Mdj2. J Biol Chem 280: 31608–31614

Morgenstern M, Stiller SB, Lübbert P, Peikert CD, Dannenmaier S, Drepper F, Weill U, Höß P, Feuerstein R, Gebert M, et al (2017) Definition of a High-Confidence Mitochondrial Proteome at Quantitative Scale. Cell Rep 19: 2836–2852

Mossmann D, Meisinger C & Vögtle FN (2012) Processing of mitochondrial presequences. Biochim Biophys Acta-Gene Regul Mech 1819: 1098–1106

Neupert W & Herrmann JM (2007) Translocation of proteins into mitochondria. Annu Rev Biochem 76: 723–749 doi:10.1146/annurev.biochem.76.052705.163409 [PREPRINT]

Nunnari J & Suomalainen A (2012) Mitochondria: In sickness and in health. Cell 148 doi:10.1016/j.cell.2012.02.035 [PREPRINT]

Pereira GC, Allen WJ, Watkins DW, Buddrus L, Noone D, Liu X, Richardson AP, Chacinska A & Collinson I (2019) A High-Resolution Luminescent Assay for Rapid and Continuous Monitoring of Protein Translocation across Biological Membranes. J Mol Biol 431: 1689

Pfanner N, Warscheid B & Wiedemann N (2019) Mitochondrial proteins: from biogenesis to functional networks. Nat Rev Mol Cell Biol 20: 267–284

Qian X, Gebert M, Höpker J, Yan M, Li J, Wiedemann N, Van Der Laan M, Pfanner N & Sha B (2011) Structural basis for the function of Tim50 in the mitochondrial presequence translocase. J Mol Biol 411: 513–519

Richter-Dennerlein R, Dennerlein S & Rehling P (2015) Integrating mitochondrial translation into the cellular context. Nat Rev Mol Cell Biol 16: 586–592

Roise D, Horvath SJ, Tomich JM, Richards JH & Schatz G (1986) A chemically synthesized pre-sequence of an imported mitochondrial protein can form an amphiphilic helix and perturb natural and artificial phospholipid bilayers. EMBO J 5: 1327–1334

Ryan MT, Voos W & Pfanner N (2001) Assaying protein import into Mitochondria. Methods Cell Biol: 189–215

Schneider HC, Berthold J, Bauer MF, Dietmeier K, Guiard B, Brunner M & Neupert W (1994) Mitochondrial Hsp70/MIM44 complex facilitates protein import. Nat 1994 3716500 371: 768–774

Schulz C, Lytovchenko O, Melin J, Chacinska A, Guiard B, Neumann P, Ficner R, Jahn O, Schmidt B & Rehling P (2011) Tim50’s presequence receptor domain is essential for signal driven transport across the TIM23 complex. J Cell Biol 195: 643

Schulz C & Rehling P (2014) Remodelling of the active presequence translocase drives motor-dependent mitochondrial protein translocation. Nat Commun 2014 51 5: 1–9

Schulz C, Schendzielorz A & Rehling P (2015) Unlocking the presequence import pathway. Trends Cell Biol 25: 265–275

Sickmann A, Reinders J, Wagner Y, Joppich C, Zahedi R, Meyer HE, Schönfisch B, Perschil I, Chacinska A, Guiard B, et al (2003) The proteome of Saccharomyces cerevisiae mitochondria. Proc Natl Acad Sci U S A 100: 13207–13212

Vögtle FN, Wortelkamp S, Zahedi RP, Becker D, Leidhold C, Gevaert K, Kellermann J, Voos W, Sickmann A, Pfanner N, et al (2009) Global analysis of the mitochondrial N-proteome identifies a processing peptidase critical for protein stability. Cell 139: 428–439

Voos W, Gambill BD, Guiard B, Pfanner N & Craig EA (1993) Presequence and mature part of preproteins strongly influence the dependence of mitochondrial protein import on heat shock protein 70 in the matrix. J Cell Biol 123: 119–126

Wiedemann N & Pfanner N (2017) Mitochondrial machineries for protein import and assembly. Annu Rev Biochem 86: 685–714

Yamamoto H, Esaki M, Kanamori T, Tamura Y, Nishikawa S ichi & Endo T (2002) Tim50 is a subunit of the TIM23 complex that links protein translocation across the outer and inner mitochondrial membranes. Cell 111: 519–528

Yamamoto H, Fukui K, Takahashi H, Kitamura S, Shiota T, Terao K, Uchida M, Esaki M, Nishikawa S-I, Yoshihisa T, et al (2009) Roles of Tom70 in Import of Presequence-containing Mitochondrial Proteins * □ S.

Yamano K, Yatsukawa Y-I, Esaki M, Aiken Hobbs AE, Jensen RE & Endo T (2007) Tom20 and Tom22 Share the Common Signal Recognition Pathway in Mitochondrial Protein Import *.

